# Demographic consequences of an extreme heatwave are mitigated by spatial heterogeneity in an annual monkeyflower

**DOI:** 10.1101/2023.05.23.541340

**Authors:** Laura M. McDonald, Anna Scharnagl, Andrea K. Turcu, Courtney M. Patterson, Nicholas J. Kooyers

**Affiliations:** Department of Biology, University of Louisiana, Lafayette, LA 70503 USA; Department of Integrative Biology, University of California, Berkeley, CA 94720 USA

**Keywords:** phenology, global change ecology, evolutionary ecology, natural selection, common garden experiment, adaptation lag

## Abstract

Heatwaves are becoming more frequent and intense with climate change, but the demographic and evolutionary consequences of heatwaves are rarely investigated in herbaceous plant species. We examine the consequences of a short but extreme heatwave in Oregon populations of the common yellow monkeyflower (*Mimulus guttatus*) by leveraging a common garden experiment planted with range-wide populations and observational studies of eleven local populations. In the common garden, 89% of seedlings died during the heatwave including >96% of seedlings from geographically-local populations. Some populations from hotter and drier environments had higher fitness, however others from comparable environments performed poorly. Observational studies of local natural populations drastically differed in the consequences of the heatwave - one population was completely extirpated and nearly half had a >50% decrease in fitness. However, a few populations had *greater* fitness during the heatwave year. Differences in mortality corresponded to the impact of the heatwave on soil moisture – retention of soil moisture throughout the heatwave led to greater survivorship. Our results suggest that not all populations experience the same intensity or degree of mortality during extreme events and such heterogeneity could be important for genetic rescue or to facilitate the distribution of adaptive variants throughout the region.

## Introduction

Climate change is not only causing gradual increases in global mean temperatures but is also causing higher levels of variation in temperature and precipitation in specific areas across the world [1]. This increase in variation suggests extreme climatic events, such as droughts, floods, and heatwaves, will become more common and more severe in certain locations [1–4]. While there is often time for species to disperse to areas with more optimal conditions during prolonged extreme events [5], pulses of extreme climate conditions can be challenging for organisms with limited movement, potentially causing severe mortality with lasting demographic consequences or even extirpation [6–8]. Such extreme climatic events in natural populations are challenging to study because these events are rare by definition, occur unpredictably, are short-lived, and often require background information or data on a specific species collected before the event to address meaningful biological questions [9].

Heatwaves, defined as three or more consecutive days where the temperature is greater than the 90^th^ percentile for a given location and time of year [10], have increased dramatically over the last century [11,12] and are predicted to further increase in frequency, duration and intensity in the coming century [12–14]. Such heatwaves and associated water-availability stress have been linked to mass mortality events in natural populations [8,15,16] and substantial loss of yield or even complete crop failure in agricultural systems [17]. However, there are relatively few studies of heatwaves documented in natural populations, especially in herbaceous plant populations, that examine how these events impact immediate population fitness and long-term population dynamics, but see [18–20]. This data is critical for determining extirpation risks for populations and future evolutionary responses [21].

While native populations may struggle with extreme conditions caused by heatwaves, geographically-distant populations that have historically experienced hotter and/or drier conditions may be better adapted to such conditions. Indeed, experimental studies from crops and model systems including *Oryza*, *Zea* and *Arabidopsis* indicate there is substantial genetic variation in escape, avoidance and tolerance to heat stress within species [22,23] and populations that experience heat stress more often are better adapted to it [24,25]. Additionally, adaptation lags, where plants originating in populations with historically geographically-distant populations are better adapted than the native population to a site because of shifting climate, have been observed frequently in the context of variation in annual climates rather than in extreme short-term climatic events [26–28].

Alternatively, there are a number of reasons why populations from hotter and drier regions may *not* produce a fitness advantage over native populations during a heatwave. Historically hotter and drier populations may not have evolved resistance mechanisms sufficient to withstand extreme and rapid heatwaves. That is, optimal physiological performance at a higher temperature is not necessarily the same agent of selection as survival during short term heat shock or performance within strongly fluctuating environments. [29]. Even if the population from the hotter/drier climate has a fitness advantage during an extreme event, it does not necessarily indicate that the associated phenotypes or allelic variation would increase in frequency within the native site. Distant populations may not be well adapted to other key abiotic and biotic selective agents within a native site [30]. This maladaptation could manifest as a tradeoff to heat resistance where populations with high survivorship during the heatwave may also have lower fecundity at the novel site relative to the native population.

In cases where heatwaves cause population declines or extirpation, natural levels of gene flow between geographically proximate populations may allow recolonization of populations that experience intense mortality or provide an influx of genetic variation (i.e. ‘genetic rescue’) [31,32]. This could be particularly important in environments with patchy habitats or in those that occur across steep environmental gradients, as geographically-close populations may not experience as extreme conditions as the focal population [33]. Assessing how nearby populations perform during an extreme event and the factors that cause heterogeneity in fitness may be as important as examining a focal population as these populations could provide an influx of individuals and genetic diversity to the focal population.

In this study, we examine how populations of a model species for ecological genetics, the common yellow monkeyflower (*Mimulus guttatus; syn. Erythranthe guttata*), perform during an extreme heatwave. Annual populations of *M. guttatus* occur throughout western North America in inland areas with ephemeral water supplies such as rock walls, seepy meadows and flood plains [34]. Annual plants germinate during spring rains or snow melt and senesce after producing seed during dry summers. The timing and length of the growing season varies dramatically across the range and there is considerable variation in the climates different populations experience [35]. Annual *M. guttatus* exhibits a wide range of phenotypic variation that allows for local adaptation to divergent environments [27,36–38] and also has some of the highest levels of standing genetic variation across plant species [39–42]. These studies suggest the genetic and phenotypic variation necessary to respond to selection due to an extreme event is likely present somewhere across this wide range.

Although *M. guttatus* is a common species unlikely to be threatened with extinction due to climate change, high elevation populations in the Central Oregon Cascades are at risk. These populations have the shortest growing season of all known annual *M. guttatus* populations, lasting only from early June to mid-July [35]. There is significant year-to-year variation in environmental conditions that can shift the growing season up to a month earlier [27,38]. While populations maintain extremely high levels of polymorphism due to temporally fluctuating selection and fine-grain heterogeneity in water availability within populations, these populations are experiencing the extremes of the historical climatic normal [38,43,44] and adaptation lags to changing conditions have already been documented [27].

Here we investigate how *M. guttatus* populations from throughout the range and within local populations in the Central Oregon Cascades perform during an extreme, but short 8-day heatwave at the beginning of the growing season. Specifically, we use a common garden experiment to compare fitness from populations throughout the range of annual *M. guttatus* and we collect phenology and fitness data from nearby populations to better understand the metapopulation-wide consequences of an extreme heatwave. We use these data to address the following questions: 1) How do local populations perform relative to more geographically-distant populations during an extreme event? 2) Do extreme weather events favor populations whose historical environment more closely matches the extreme event? 3) Are the consequences of heatwaves homogenous across a metapopulation?, and 4) If not, is heterogeneity predictable by variation in environmental characteristics between sites? We find that native populations performed poorly in our common garden, but some distant populations that historically encounter extreme heat more frequently have far higher fitness. Local populations also show considerable variation in responses to the heatwave with some populations avoiding negative fitness consequences entirely.

## Materials and Methods

### Documenting an extreme event

During the second week of June 2019, we observed abnormally high temperatures and rapid dry down following snowmelt in our long-term common garden and observation sites in the central Oregon cascades. We took advantage of our existing infrastructure and ongoing experiments to examine the impact of this heatwave on *M. guttatus* populations. We quantified the magnitude of this heatwave by comparing climate in 2019 to historic averages. We downloaded monthly averages of minimum, average, and maximum temperatures as well as average precipitation and precipitation as snow for each year from 1980-2019 from ClimateWNA [45] for the Browder Ridge, Oregon common garden site (44.37348, -122.13055, Elev. = 1246m asl). For more precise and frequent temperature observations during the growing season, daily summary data from 1980-2019 was downloaded for the closest NOAA weather station (Santiam Jct.: 44.44, -121.95, Elev. = 1140m asl, 16km from the garden) from NOAA’s National Climatic Data Center.

### Common Garden Study

To examine how the heatwave influenced relative patterns of adaptation, we leveraged a common garden field experiment that contained outbred lines from 11 populations spanning much of the range of annual *M. guttatus* at the Browder Ridge site (Fig. 1A). Each outbred line was derived from seeds collected from 5-8 maternal lines (Ave. 7.1). Latitude, longitude and elevation of each site was taken at time of collection and used to acquire climatic norms (1960-1990) for each population from ClimateWNA as well as haversine distances from each population to the Browder Ridge common garden. To generate outbred lines, maternal lines were grown for a single generation in a common garden greenhouse environment and crossed to another maternal line from the same population. Each maternal line acted as a pollen recipient in one cross and a pollen donor in a second cross.

**Fig. 1.**
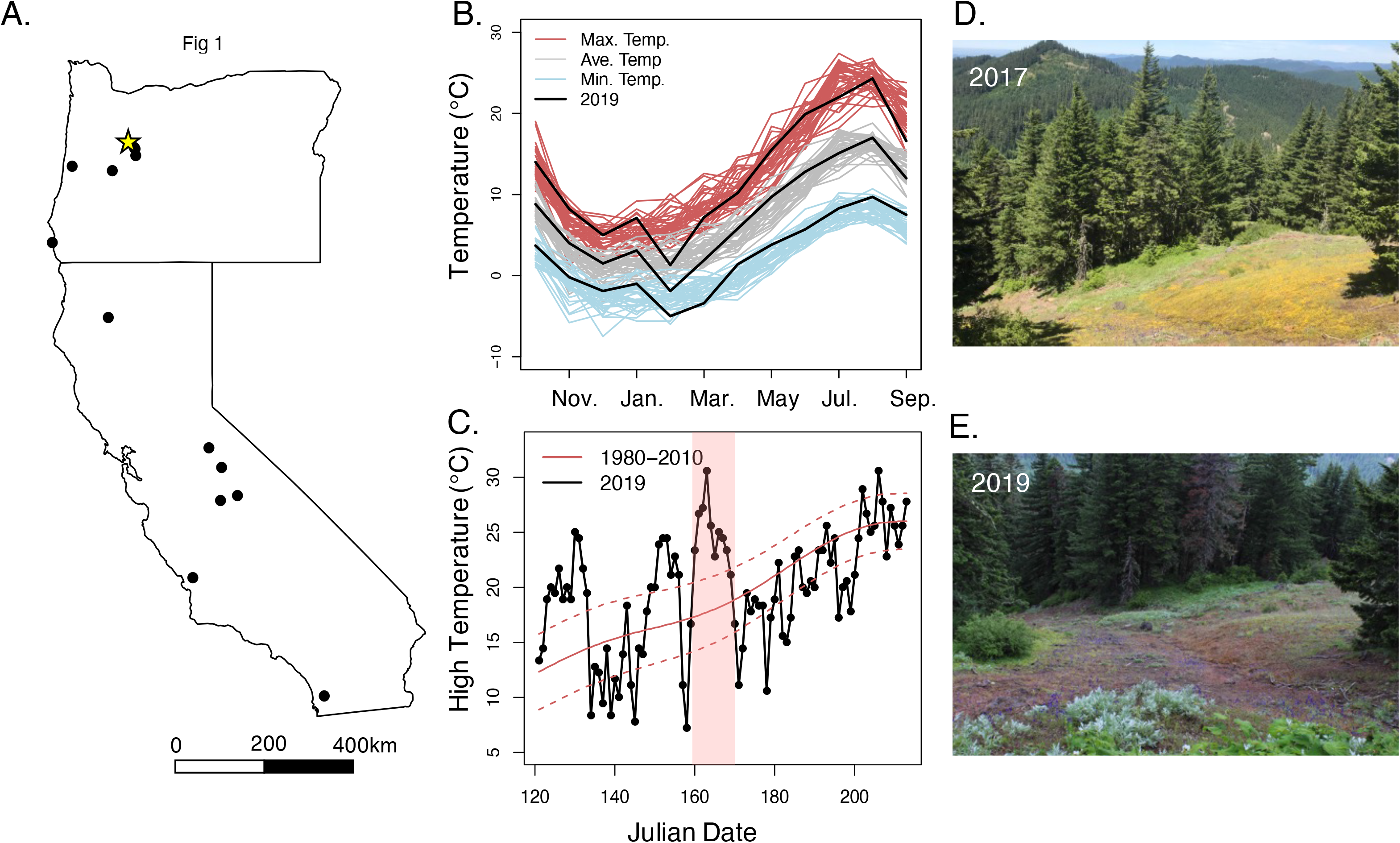
Locations of populations and description of weather patterns during the 2019 growing season. A) Map of the populations used within the common garden experiments (black dots) as well as the location of the Broader Ridge common garden site (yellow star). B) Annual climate patterns from 1980 - 2019 constructed from monthly averages of minimum (blue), average (gray) and maximum temperature (red). Black lines are data from the 2019 growing season. C) Maximum daily temperatures during the 2019 growing season (black). Solid red line represents the historical average maximum temperatures (1980-2010) for each day and dashed lines represent 99% confidence intervals. Vertical red bar indicates days of extreme heatwave. D, E) Photographs taken in the exact same location at the nearby Iron Mountain population of *M. guttatus* during 2017 and 2019.

We initiated the common garden by planting outbred line seeds in 2.5” pots filled with Sunshine #1 (Sun Gro Horticulture; Massachusetts, USA) in unperforated 10”x 20” flats. We covered the flats with clear humidity domes, and cold-stratified the seeds in the dark at 4°C. After seven days of stratification, we moved the flats to the University of Oregon greenhouse into ambient light and temperature conditions. Plants were misted and germination was recorded daily. Following seven days in the greenhouse, we removed humidity domes and plants were bottom-watered as needed. After 14 days in the greenhouse, we randomized all pots into twelve blocks and transplanted seedlings directly into the field site. Microsite variation in water availability is high at this site with a natural population of *M. guttatus* spanning areas that dry out at different rates. We planted the blocks in locations that span this variation. Loss due to transplant shock has been minimal at this field site in past years [27,38,46]. Timing matched the phenology of local populations. That is, all native plants were vegetative rosettes within a few weeks of germination.

We surveyed survival and flowering for each plant every other day. Survival was defined as having any living green tissue in leaves, stem or meristem (i.e. active chloroplast activity). No plant that was recorded as dead appeared later in the growing season as alive. To assess fecundity, we counted the number of flowers, and collected all mature calyxes and counted the seeds they contained. Below we report ‘inclusive number of flowers’ as total number of flowers where plants that did not survive or flower counted as zeros and ‘inclusive number of seeds’ as number of seeds produced where plants that did not produce seeds counted as zeros. We report both metrics because number of flowers better includes fitness contributions through male fecundity while number of seeds better represents female fecundity.

### Assessing differences in fitness between populations in the common garden

We determined how the native population performed relative to other populations throughout the *M. guttatus* range using linear mixed models and generalized linear mixed models (LMMs and GLMMs) implemented with the *lmer()* and *glmer()* functions in the *lme4* v1.1-27.1 package [47] in R v4.1.1 (Institute for Statistic Computing; Vienna, Austria). Separate univariate models were constructed for five different fitness components (survival to flowering, number of flowers, number of seeds, inclusive number of flowers, and inclusive number of seeds) as the response variable. Number of seeds and inclusive number of seeds were both log-transformed to improve the fit of the below models. Each model had population as a fixed factor and line as a random factor. Additionally, variation among blocks in the garden was included as a random factor in both models. The GLMM assessing survival to flowering had a binomial error distribution with a logit link while LMMs were used for number of flowers, number of seeds, inclusive number of flowers and inclusive number of seeds (GLMMs using a Poisson family and logit link had nearly identical results; Table S1). Statistical significance of population was assessed via ANOVA using the *Anova()* function in the *car* v3.0-12 package [48]. We compared the native population to all other populations by calculating line means for each variable and conducting Dunnett’s tests with BR1 as the focal population. Dunnett’s tests were implemented using the *DescTools* v0.99.44 package. We note that our estimates of absolute fitness are likely elevated from natural populations as we transplanted seedlings to limit initial mortality.

We also investigated potential tradeoffs between survival of the heatwave and fecundity by comparing whether lines that had a higher chance of surviving the heatwave were more likely to have higher fecundity. We constructed linear models to examine the association between probability of surviving until flowering for a given maternal line and the average number of flowers or average number of seeds that the maternal line produced. Models were implemented using the *lm()* function and number of seeds was log transformed as above. Additional models with a logit transformation of probability of surviving until flowering produced qualitatively identical results. A negative correlation between probability of survival and fecundity is indicative of a tradeoff.

### Fitness-historical environment associations

We examined whether historic climatic differences among populations were associated with differences in fitness using a univariate GLMM approach with fitness variables as the response variables in independent models. All models included block, population and line nested within population as random variables and the environmental variable as a fixed factor. For each fitness variable, we ran separate GLMM for seven different environmental variables including: geographic distance to the Browder Ridge common garden, mean annual temperature, annual heat moisture index, growing season start date, precipitation as snow, variance in spring maximum temperature and variance in summer maximum temperature. Variances were calculated from maximum spring and summer temperatures from 1960-2021 extracted from ClimateWNA. These factors were chosen because they have all been identified as potential agents of selection for *M. guttatus* in past studies [27,35]. Error distributions, links and transformations are the same as described above. Statistical significance was determined by ANOVA as above.

### Impact of the heatwave on natural populations

We selected eleven natural populations distributed across an elevational gradient of ∼600m in a 10km^2^ region of Browder Ridge to examine variation in soil moisture, survivorship, and phenology across the 2019 growing season (Fig. S1). In each population, we surveyed two 0.25m^2^ sampling grids (50cm x 50cm) every 7-14 days from snowmelt until population senescence. Each visit, we counted the number of vegetative, flowering, and senesced plants as well as measuring volumetric water content with a SM150T soil moisture sensor (Dynamax; Texas, USA). Grid locations were chosen within sites to both encompass the natural variation within site while only using areas with high concentration of *M. guttatus* seedlings.

We examined how number of individuals in each grid changed before, during and after the heatwave to assess the mortality associated with the heatwave. To examine whether differences in soil moisture were driving differences in mortality between plots, we used a LMM to model whether the amount of mortality experienced in a grid during the heatwave was associated with the volumetric soil water content before the heatwave. Population was treated as a random factor in these models. Statistical significance of the fixed factors was assessed with lmerTest v3.1-3 using the Satterthwaite’s degrees of freedom method [49]. We further examined variation in survivorship, phenology and volumetric water content throughout the growing season by modeling mortality, flowering and soil moisture through time and calculating summary statistics for each grid (see Supplemental Methods). Summary statistics included critical survivorship date (when 50% of plants still survived in a plot), peak flowering date, and date when VWC fell below 20% for the first time.

### Impacts of the heatwave on fecundity in natural populations

We examined the influence of the heatwave on fecundity using observational data collected at the end of the 2018 and 2019 growing seasons from twelve natural populations. Nine of these populations were the same ones as described above (Fig. S1). We chose plants in each population using two 7.5m transects running through the center of each population. The same transect locations were used in 2018 and 2019. We identified the closest plant to the survey line every 15cm along the transect and counted number of flowers and seeds for each plant. We collected 100 plants/population when possible, but did not make the full collection in every population in 2019 as three populations had very few plants producing seed (HDM, OWC, RRM). For these populations we collected <10% of the total individuals within the population. We modeled how number of flower and number of seeds changed between years and populations using a LMM with log of seed set as the response variable and Population, Year, and Population:Year interaction as independent variables. Statistical significance was determined using *Anova()* as above with a type III sum of squares. Because the interaction between population and year was significant, we examined population means to determine the direction and magnitude of differences in each individual population between years. We also evaluated whether the relative differences in flower or seed production between 2018 and 2019 that we observed between populations was associated with the environmental characteristics of the populations. We modeled associations between difference in seed production between years and each of six different environmental characteristics (elevation, mean annual temperature, mean coldest month temperature, beginning of the frost free period, precipitation as snow and climate moisture deficit) using linear regressions implemented through the lm() function. Differences in seed production between years was calculated from population averages.

## Results

### Early season heatwave decimates M. guttatus experimental garden

Climate patterns in 2019 in the Central Oregon Cascade Mountain Range closely resembled historic monthly normals for both temperature and precipitation (Fig. 1B). However, a severe heat wave occurred approximately two weeks following snow melt at the Browder Ridge field site. Data from the nearest NOAA weather station indicates max temperatures over an 8-day period were the hottest on record back to 1983 (Day 160-167; Fig. 1C) and peaked at 30.4° C on Day 163 (June 12th). The average maximum temperature during this stretch in 2019 exceeded the historic average maximum temperature by 8.5° C. In the Browder Ridge common garden, 89.0% of seedlings (438/492) died during the heatwave. There was a significant effect of block on survivorship (X^2^ = 141.1, p < 0.001). Survivorship in blocks ranged from 0 - 77.3% and only three blocks had greater than 15% survivorship. Blocks where soil dried out later in the heatwave had higher survivorship suggesting that death was due to a combination of limited water availability and heat stress (A. Scharnagl pers. obs.).

### Native populations have low relative fitness in common garden

There were significant differences in survivorship among populations planted in the common garden following the heatwave (X^2^ = 22.9, p =0.01, Table S1). Populations from locations geographically close to the common garden site with similar historical climate conditions had extremely low survivorship following the drought (Fig. 2A, Table S2). The BR1 population did not have a single individual survive the heatwave (0/21 seedlings; 0.38 km away) and the MTC population had only two individuals survive (2/37 seedlings; 15.6 km away). Instead, three more distant populations exhibited notably higher survivorship than the other populations. A nearby low elevation population (LPD, 71.2 km away) had 16.4% of seedlings survive the drought, a population from the Klamath Mountains (TAR, 396 km away) had 17.2% survival, and a population from the foothills of the Sierra Nevada Mountains (BEL, 839 km away) had 24.2% survival.

**Fig. 2:**
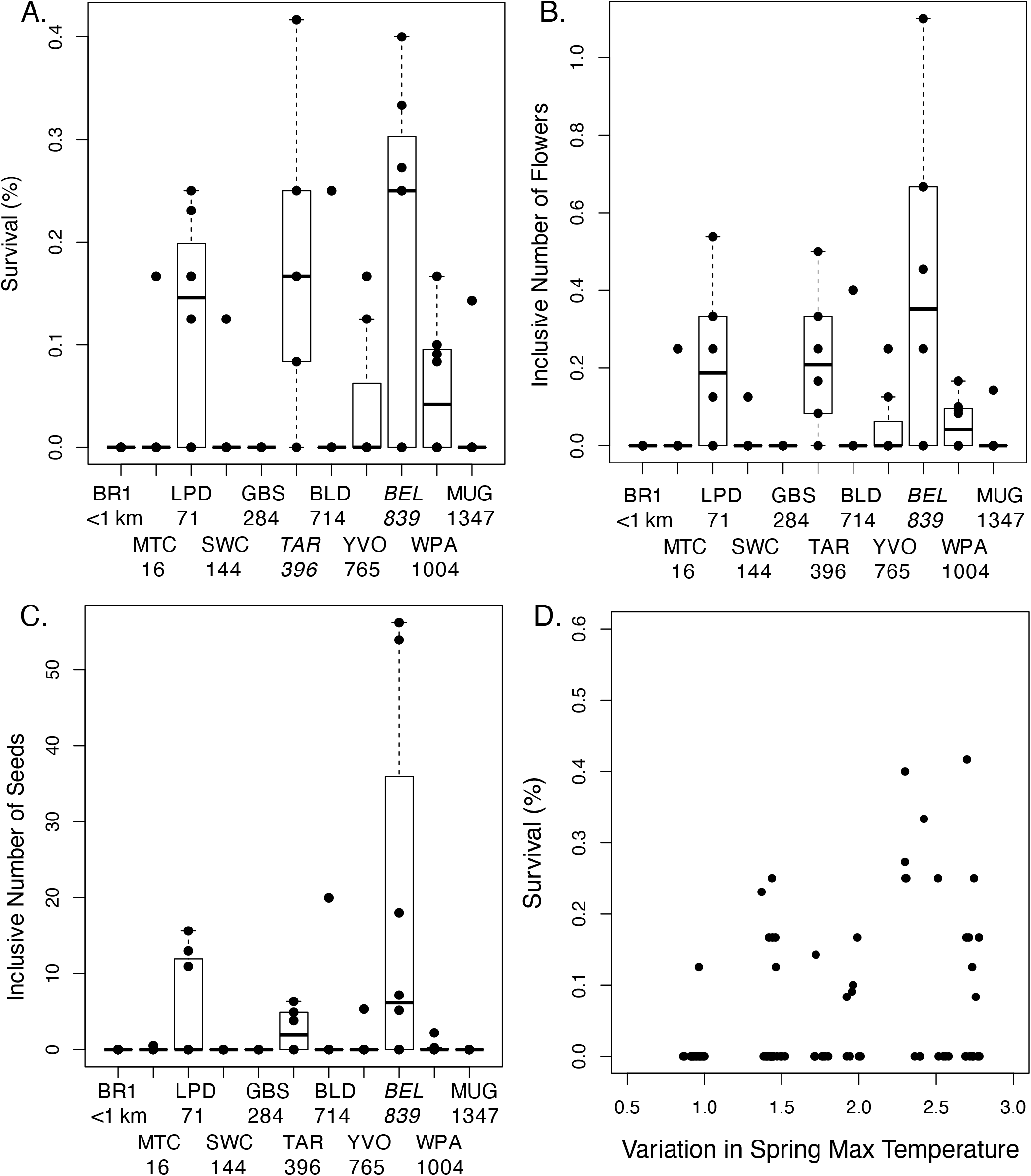
Variation in fitness between populations within the common garden experiment. Survivorship to flowering (A), inclusive number of flowers (B) and inclusive number of seeds (C) for each population within the common garden experiment. Points represent line averages. Values below population abbreviations are the distance from the Browder Ridge common garden site in kilometers. Italicized populations names are the populations that significantly differ in survivorship from the BR1 population at α= 0.05 in a Dunnett’s test. (D) Association between survival to flowering in the common garden and the source population’s variance in spring maximum temperature. Points represent the percentage individuals that survived to flowering within a given line. Points are slightly jittered in order to better see the number of lines per population with no survival.

Following the heatwave, there were no significant differences in fecundity among populations (Number of Flowers: F8,42.6 = 1.8, p = 0.10; Number of Seeds: F6,10.4 = 1.4, p = 0.30, Table S1) and metrics of fitness that include both survival and fecundity closely resembled survival models, with significant variation among populations (Inclusive Number of Flowers: X^2^ = 47.0, p < 0.0001; Inclusive Number of Seeds: X^2^ = 40.1, p = 0.0002; Fig. 2). Of the four populations that produced >1 seed/plant on average (i.e. an approximation of replacement level), the closest population to the BR common garden site was a low elevation Oregon site (LPD) located 71.2km away and the other three populations were from California. There was no evidence of a tradeoff between ability to survive the heatwave and fecundity following the heatwave. Lines that had higher survival during the drought produced more flowers than lines that had lower survival (r^2^ = 0.29, p = 0.006; Fig. S2), but there was no relationship between viability and the number of seeds produced (r^2^ = 0.07, p = 0.28, Fig. S2). Together these results suggest that local populations are less likely to survive and do not have a fecundity advantage over geographically distant populations following an early season heatwave.

### Populations from more arid areas do not necessarily perform better

Although there were clear differences among populations in both survivorship following the heatwave and inclusive number of flowers and seeds, these differences are not tightly associated with the historic environments of the populations (Fig. 2D, S3; Table S3). The only variable even marginally associated with any fitness metric was variation in spring maximum temperature with survival (X^2^ = 3.6, p = 0.06; Fig. 2D). The three populations that best survived the heatwave were from notably warmer and more arid climates than Browder Ridge but there are other populations from similarly warm/arid climates that had very low survivorship (Fig. S3). Likewise, neither inclusive number of flowers nor seeds was associated with any geographic or historical climate predictor (Table S3).

### Widespread but variable mortality across a metapopulation

To better understand how this extreme heatwave could impact *M. guttatus* populations across a relatively small geographic region, we followed the survivorship and phenology of 11 populations (two 0.25m^2^ sampling grids per population) at locations across Browder Ridge. Indeed, the heatwave *did* cause mortality for every population surveyed except one (MBR). Number of individuals in 23 of 24 grids declined during the heatwave (Fig. 3A). While the number of individuals per grid increased by 10% on average in the week prior to the heatwave, the average number of individuals in a grid declined by 50.4% (sd: 30.96%) between the pre and post-heatwave sampling points (Table S4). One population (HDM) had no individuals from either grid survive the heatwave. Other populations had extreme differences in survivorship between grids within a single population. For instance, one grid in BR1 had only 2% mortality during the heatwave while the other grid, located only ∼1.5m away, had 53.4% mortality. There was no relationship between elevation or other coarse environmental factors and survivorship in each grid during the heatwave (Table S5). This lack of a pattern suggests small-scale microclimatic variation found within a population is most predictive of survivorship.

**Fig. 3.**
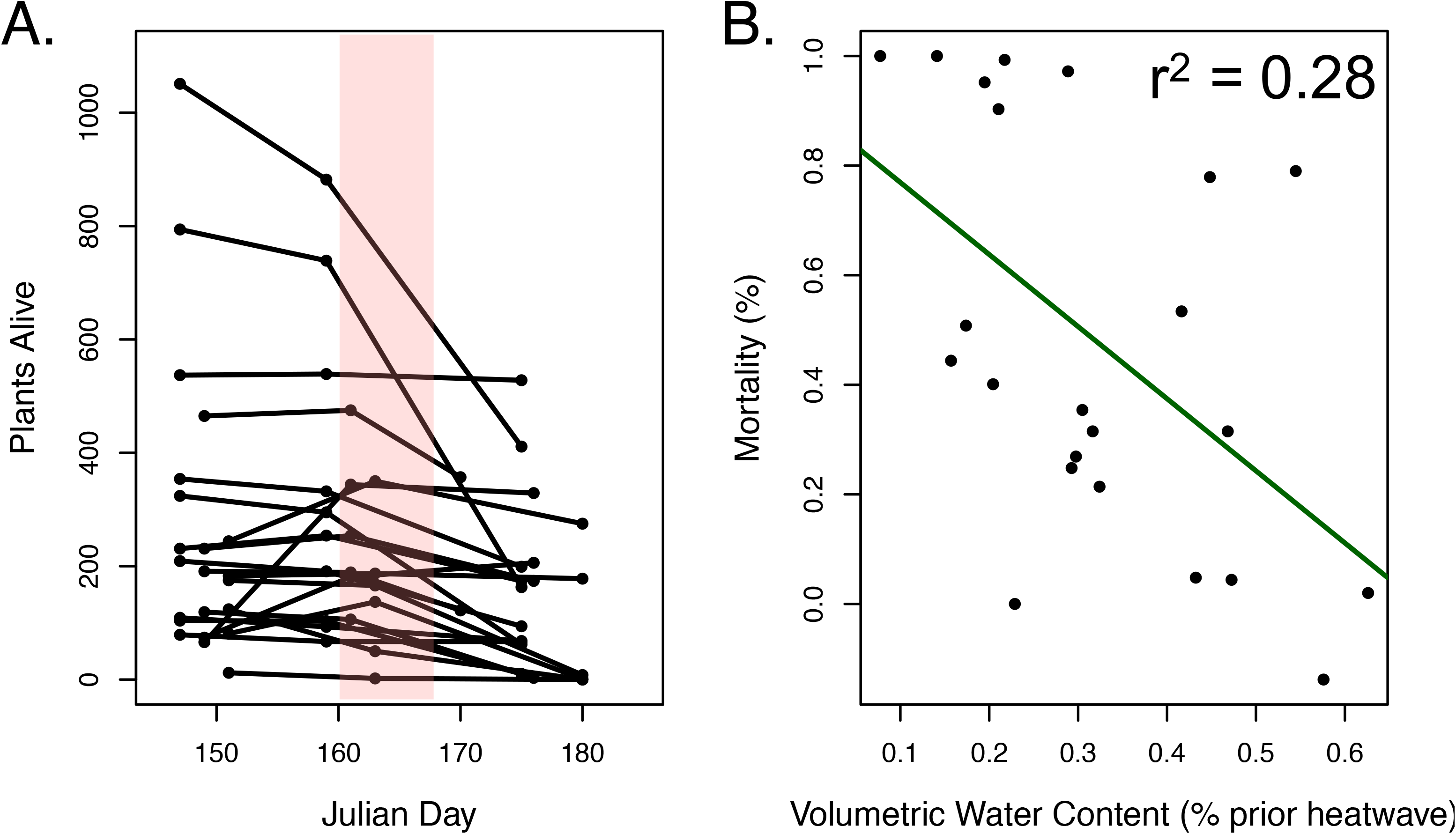
Responses across natural populations to the extreme heatwave. (A) Plot depicts the number of plants alive in each grid surveyed before, during and after the heatwave (the heatwave occurred from day 160 to 167). (B) The relationship between mortality and soil moisture prior to the heatwave in each grid. Solid line is the regression line of best fit.

### Soil moisture is associated with survivorship

Since *M. guttatus* populations are characterized by having ephemeral supplies of water, we examined how soil moisture, survivorship and phenology varied within and between natural populations as a potential key factor for understanding responses to the heatwave. Nearly all sampling grids dropped below 20% VWC during the heatwave, although several populations, particularly at higher elevations, returned above 20% later in the growing season (Fig. S4).

There was a strong association between soil moisture before the heatwave and the mortality during the heatwave where grids with lower VWC before the heatwave had higher mortality (X^2^ = 9.1, p = 0.003; Fig. 3B) – that is grids that dried down earlier had plants die earlier. Peak flowering occurred after critical survivorship dates in 76% of grids indicating that most plants died before flowering. There were substantial differences in peak flowering among populations with peak flowering occurring later in populations with later dry down dates (Fig. S5). Combined, these data suggest that survival and phenology in these populations are strongly associated with variation in soil moisture rather than the actual heat stress associated with the heatwave.

### Heterogeneous responses of the heatwave across the metapopulation

We compared fecundity measures from 12 populations between 2018 (relatively normal year) and 2019 (extreme heatwave early in growing season). There was a significant population by year effect on both number of flowers (F7, 1484 = 12.3, p < 0.0001) and number of seeds (F7, 1476 = 21.3, p < 0.0001). The majority of natural populations had lower fecundity during the year of the heatwave (2019). In the populations that did worse in 2019 than 2018, the number of flowers and number of seeds were reduced by an average of 36% and 41%, respectively (Fig. 4, Table S4). However, some populations had higher fecundity during the heatwave year. Two populations produced more flowers in 2019 than 2018 (Fig. 4; FIR and SEC) and four populations produced more seed (Fig. 4; FIR, SEC, OWC, SMG). Differences in fecundity between 2018 and 2019 were not associated with elevation, distance between populations or any other environmental correlate that we examined (Table S6). These data suggest that although the entire metapopulation experienced a heatwave, there was a large variation in how the heatwave impacted monkeyflower populations that is not predictable by the historic climate of a population.

**Fig. 4.**
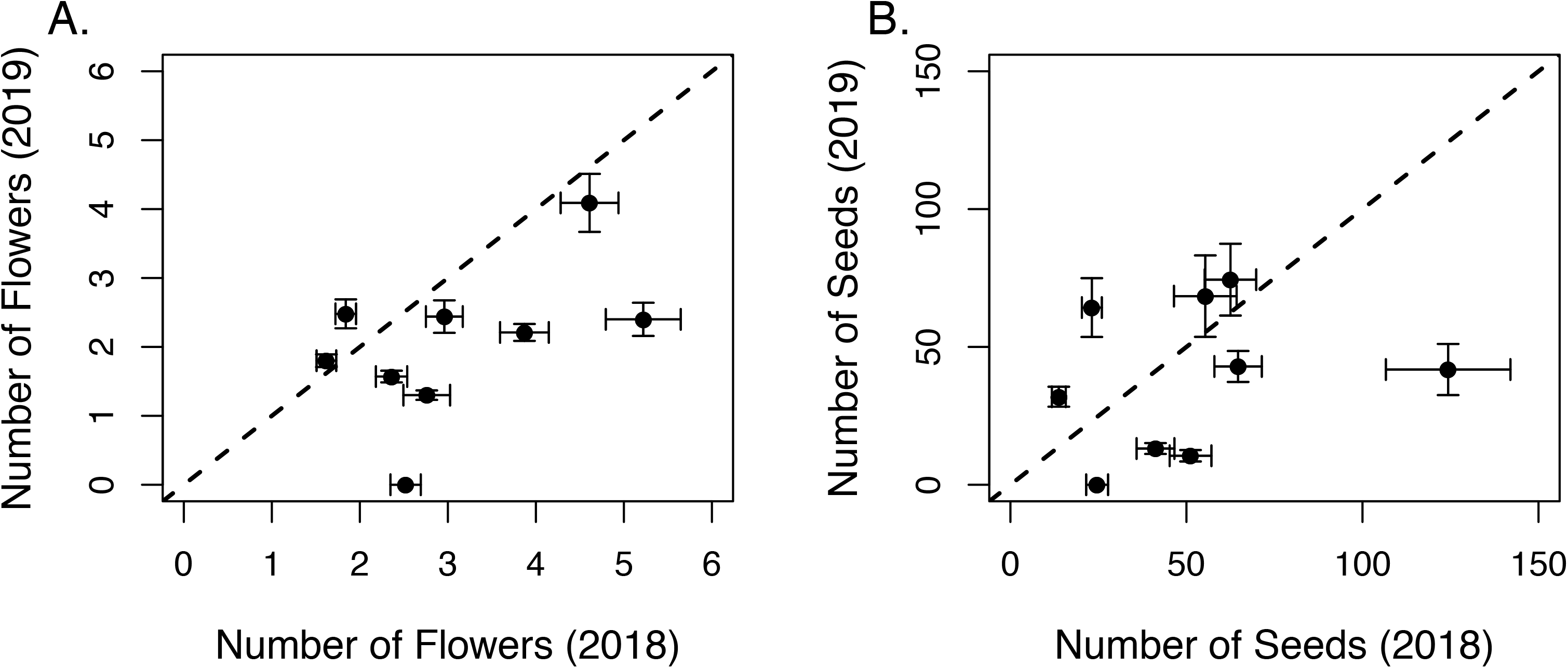
Comparisons of fitness in Browder Ridge monkeyflower populations in 2018 vs 2019. Scatterplots depicting number of flowers (A) and number of seed (B) in 2018 vs 2019. Error bars represent standard error.

## Discussion

The demographic consequences of extreme events, such as heatwaves, are rarely explored, but such results are increasingly important in a changing climate. Our study provides a comprehensive examination of the heterogenous consequences of a short but extreme heatwave for local and geographically distant populations of *M. guttatus*. Specifically, we document that Central Oregon populations experienced an extreme heatwave where high temperatures exceeded previous records. Plants from populations located near our common garden site were not well adapted to survive this event and those that did produced very few seeds. However, several of the more distant populations were better able to survive and produce seeds. While these surviving populations did come from areas with greater variation in spring max temperature or historically lower annual heat-moisture indexes, other populations from similar areas did not have higher fitness. The majority of natural populations near our common garden had lower fitness in the year of the heatwave relative to the previous year and one population became extirpated. However, some native populations did not experience the same degree of mortality and this variation was strongly associated with soil moisture levels. Together these results suggest that, even though some populations in the metapopulation could exhibit substantial decline or even extirpation, other nearby populations may not experience conditions as harsh and could act as source populations for highly-impacted populations. Below, we discuss the implications of our results and compare them to other studies of population dynamics following extreme events.

### Mortality within a heatwave

Our results indicate mortality during a heatwave can be extreme and may have long lasting consequences for a population. While our heatwave was relatively short, we observed high mortality within our common garden experiment (∼90% of plants died) as the heatwave occurred just as experimental plants were finishing establishing (transplanted 7 days prior to start of heatwave). While this could be considered mortality due to establishment shock, we suggest this is relatively unlikely given our observations in nearby natural populations. That is eleven of the twelve natural populations had drastic declines in soil moisture and high mortality during this period (Fig. 3A). Rates of mortality in natural populations pre- and post-heatwave averaged 50.3% (SD 36.3%, Range -4.4 – 100%). These rates are higher than many other studies documenting natural heatwaves and/or droughts in plant populations despite being a very short heatwave in a relatively normal year [19,20,50,51] but see [52]. For instance, estimates of mortality from 17 plant species during a two-year heatwave/drought in Western Australia averaged 26.0% mortality (SD 24.0%, Range 0 – 71%) [8]. However, the majority of previous studies of heatwaves may not be comparable as they have largely focused on longer events that are also associated with drought [53] or examined longer-lived species [8,54]. Basic data on survival of herbaceous plants to extreme conditions in nature is a necessity for predicting future plant population dynamics but is currently in short supply.

### Historical aridity does not accurately predict heatwave survival

While mortality was high within both our common garden and natural populations, the fact that a minority of plants did survive and reproduce suggests there may be traits that facilitate survival. We expect populations from areas with historic climates that more closely match the extreme event to have higher fitness than native populations that rarely encounter such conditions, i.e. an adaptation lag [26–28]. Alternatively, these distant and potentially preadapted populations could perform as badly or even worse than native populations because the distant populations may be poorly adapted to other agents of selection within the native population [55,56] or may have a tolerance strategy not well suited to short but extreme events [57]. Our results fall somewhere between these two extremes – local populations had high mortality and low fitness, and some distant populations from hotter drier areas had higher fitness (Fig. 2, Table S2). However, several populations that we assumed would do well under heat stressed conditions (i.e. populations from southern California and the high elevations in the Sierra Nevada) also had very low fitness.

A key question is why some populations that are historically warmer and drier did not have improved survival of the heatwave. We suggest that variation in traits among the populations likely provides some explanation. For instance, one of the populations with relatively high survival during the drought, LPD, is a low elevation site located only 71.2km from the common garden site. This site does not typically experience the higher temperatures and aridity of most of the California populations, but may be better adapted to the Cascades through having more similar growing season timing or biotic interactions. Indeed, low elevation Cascades plants do have more similar photoperiod requirements for flowering and chemical defense arsenals to high elevation Cascades populations than the California populations used within this study [35,55,58]. Notably there are also large differences in magnitude and traits involved in plastic responses to dry down conditions for the different populations used within this experiment, including differences in responses among California populations that could explain differential mortality. Future work should aim to link variation in trait variation and physiology to fitness consequences during extreme events.

One clear conclusion from our results is that the relative fitness patterns found in the heatwave-associated growing season (2019) do not match previous patterns found at the same location. We conducted a similar common garden experiment at the same site with many of the same populations in 2014 [27]. The 2014 growing season was one of the earliest growing seasons on record and, as in the 2019 growing season, some populations from California had higher fitness than native Oregon populations. However, a low elevation Sierra Nevada population that performed very poorly in 2014 (BEL) had the highest fitness of any population in 2019 and the high elevation Sierra Nevada populations from that did well in 2014 had relatively modest fitness in the 2019 season (Fig. 2; Table S2). This difference is likely due to the nuanced changes in selection pressures between years. In 2014, abnormally high spring temperatures led to the growing season starting weeks earlier than normal, but precipitation was relatively normal throughout the year. In 2019, spring temperatures and snowfall were near average leading to a relatively normal growing season start date prior to the early season heatwave. Thus, in 2014, populations able to take advantage of the earlier than normal growing season were presumably favored while, in 2019, plants able to survive a high heat and low water climatic event as seedlings were likely favored. Temporal heterogeneity in selection pressures in these populations has been widely documented previously [38,43], but this study demonstrates the fluctuations in environment are substantial enough to enable populations with the lowest relative fitness in one year to have the highest relative fitness another year. These results suggest predicting a population’s tolerance to an extreme event is not as simple as examining historic environmental variation in the selective pressure under question.

### Metapopulations experience variation in the consequences of extreme events

Here we report that the heatwave had severe consequences on natural populations near the common garden site including complete mortality within one population and >50% mortality prior to flowering in several other populations (Fig. 3A). Our results suggest that the most important factor for predicting mortality during the heatwave was the amount of nearby soil moisture present prior to the heatwave (Fig. 3B). This suggests that water rather than heat may have been the limiting factor during this heatwave. The consequences of this heatwave extended to differences in fecundity across most populations, with lower numbers of flowers and seeds produced per plant than in a more normal year (Fig. 4). We hypothesize that the link between mortality and fecundity stems from either the delayed growth of seedlings following the heatwave or the selective mortality of smaller seedlings or rapidly reproducing plants during the heatwave. These hypotheses stem from observations of delayed phenology relative to other years within these populations (Fig. S4; N. Kooyers pers. obs.). However, we cannot rule out altered interactions with pollinators or herbivores [59].

While the amount of mortality seems like a relatively dire result, populations seldom exist in isolation and nearby populations may influence focal population dynamics both during and following an extreme event. Importantly, nearby populations may not experience the same severe selective conditions that a focal population receives and thus can act as a source for new migrants and replenishing genetic variation following an extreme event through dispersal or gene flow [32,60]. We suggest the heterogeneity we observe in mortality within and between populations mitigates the potential for complete extinction within the metapopulation due a single extreme event and could allow for recolonization of extirpated sites. Gene flow between populations is likely high as there is very limited population structure between these populations [41]. Recolonization has been frequently observed in other monkeyflower populations. For instance, over 25 years observing 39 perennial populations of *M. guttatus* in the Wasatch mountains of Utah, there were 54 population disappearances and 34 reappearances the likely stemmed from seeds dispersing down rivers from upstream populations [61].

However, in our system, gene flow from other populations may not be necessary for an influx of individuals or genetic variation as much of the variation in heatwave-associated mortality was contained between grids within populations. The most important factor in predicting mortality was the soil moisture prior to the heatwave (Fig. 3B), and there was substantial variation within populations in soil moisture. The highest elevation population (HOV) is a perfect example of this heterogeneity within a population: while one grid still had trace amounts of snow during the beginning of the heatwave and had nearly no mortality (4.8% dead), the other grid dried down during the heatwave and had nearly complete mortality (99.3% dead). Thus, despite very high mortality within different microclimates within a population, dispersal within populations could allow for rapid recovery from extreme events.

For populations that experience high mortality, an equally viable solution is the presence of a seed bank [62,63]. Seed banks in *M. guttatus* have long been hypothesized [61], and our study provides additional evidence. We followed the ‘extirpated’ population during the heatwave for the next two years (HDM). While there was no germination the following year (2020), there were a limited number of germinants in 2021 (30-40 individuals, S. Innes pers. obsv.). The remote nature of this population and number of germinants suggests that these germinants came from the seed bank rather than from dispersal. Our study highlights the difficulty in predicting species responses to extreme events. Detecting the risk of extirpation for other species requires determining how the particular event impacts the environment, how variation in the environment corresponds with variation in mortality, and accounting for reestablishment from seed bank, dispersal and gene flow.

### Long-term consequences of extreme heatwaves

Longer term survival in a changing climate may require more than just resurrection of populations from the seed bank or from genetic rescue from other nearby populations. The heterogeneity in mortality from the heatwave across populations suggests our metapopulation is a ripe environment for the rapid evolution of heat and drought-resistance strategies [64]. Such evolution to extreme events has been described in numerous other systems e.g. [65–67] and can have long-term consequences for the populations [65]. Given high levels of genetic and phenotypic variation present in our monkeyflower populations [39,41] and a high degree of fine-grain spatial and temporal heterogeneity in environmental factors that has promoted balancing selection [38], heat or drought-resistance related alleles may already be present within certain populations at low frequencies. These populations also exhibit very limited population structure which suggests that gene flow is likely high [41]. Thus, an adaptive variant that evolves in one population will likely spread to other nearby populations relatively quickly.

In conclusion, this study suggests that extreme heatwaves can cause drastic declines in native populations, but such mortality may be ameliorated by micro-environmental variation, seed banks, and potentially from genetic rescue stemming from nearby populations. While this result is optimistic, we caution that survival of a single short term extreme event is not necessarily predictive when extreme events become the normal.

## Acknowledgements

This work was possible with logistical support from Matthew Streisfeld, Susan Belcher, the University of Oregon Greenhouse, and H.J. Andrews Experimental Forest. Joshua FitzPatrick, Bryan Luke Rabalis, and Haley St. Martin provided assistance processing seeds. We thank Cheryl Friesen for advice and assistance obtaining permits. All research was permitted through the USFS within the Willamette National Forest. Funding for this research came from University of Louisiana, Lafayette and NSF grant DEB-2045643 to NJK.

