## Supplemental Figures for "Demographic consequences of an extreme heatwave are mitigated by spatial heterogeneity in an annual monkeyflower"

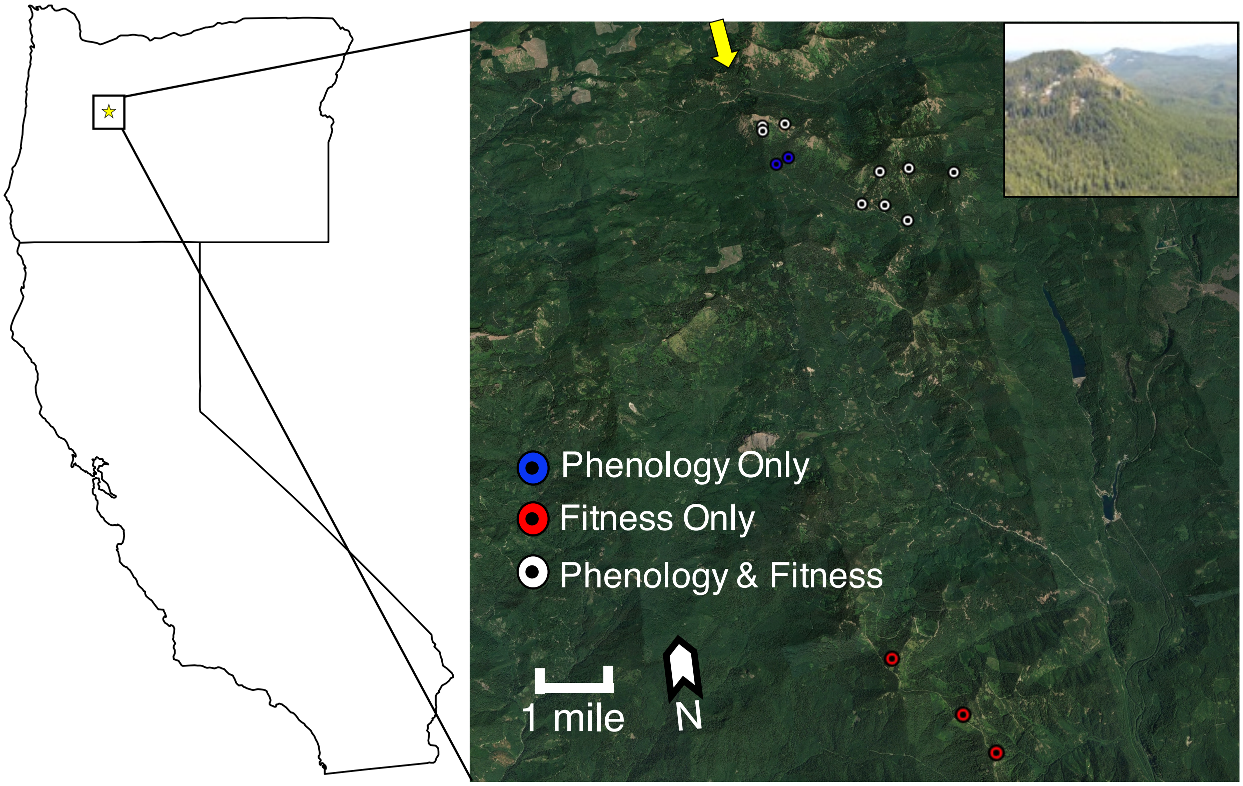


**Fig. S1:** Map of Browder Ridge monkeyflower populations. White points correspond to populations used for phenology and fitness measurements within natural populations. Red points correspond to populations used for fitness and blue to populations used only for phenology measurements. Yellow arrow reflects the location and direction that the insert photo was taken.

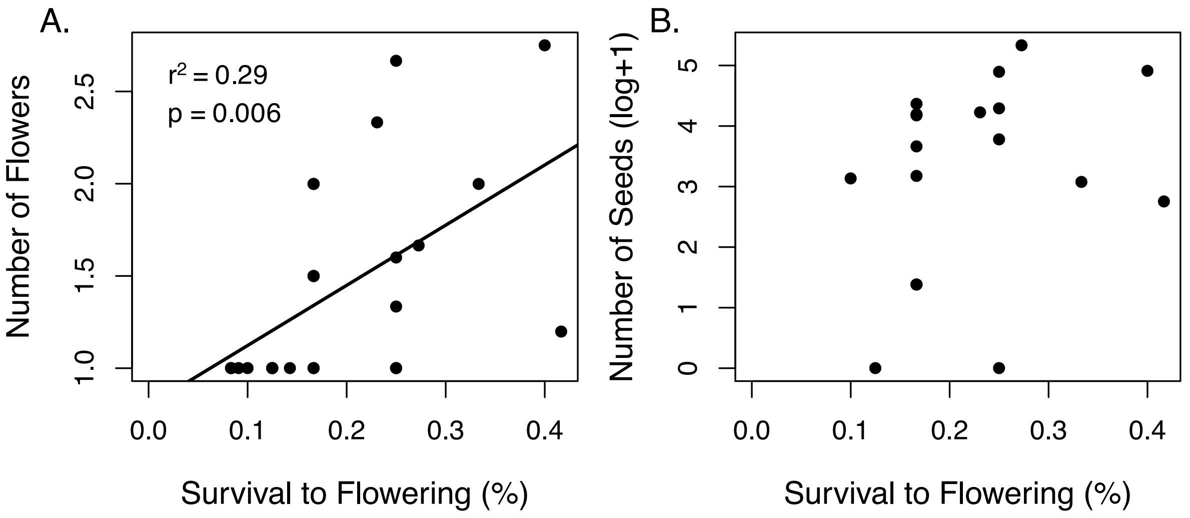


**Fig. S2** Relationships between fitness measures in the common garden study. Plots depict associations between survival to flowering and either number of flowers produced (A) or number of seeds produced (B) during the 2019 growing season. Each point represents an outbred line mean.

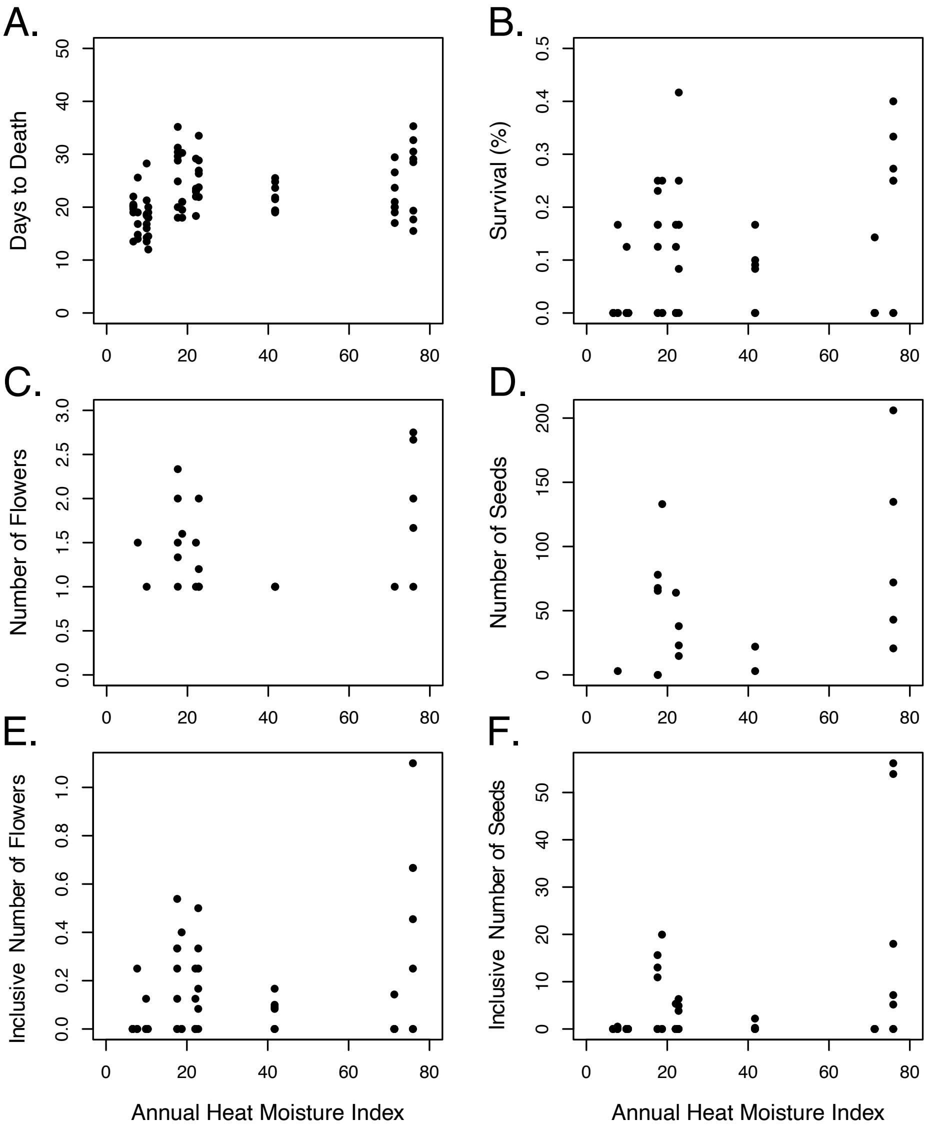


**Fig. S3**: Relationships between fitness proxies measured in the common garden and Annual Heat Moisture Indexes (AHM) from the source population locations. Points represent line averages or, in the case of survival to flowering, the percent of individuals that survived to flowering within a given line. The only significant association is between Number of Seeds and AHM (D; $X^2 = 5.9$, $p = 0.01$).

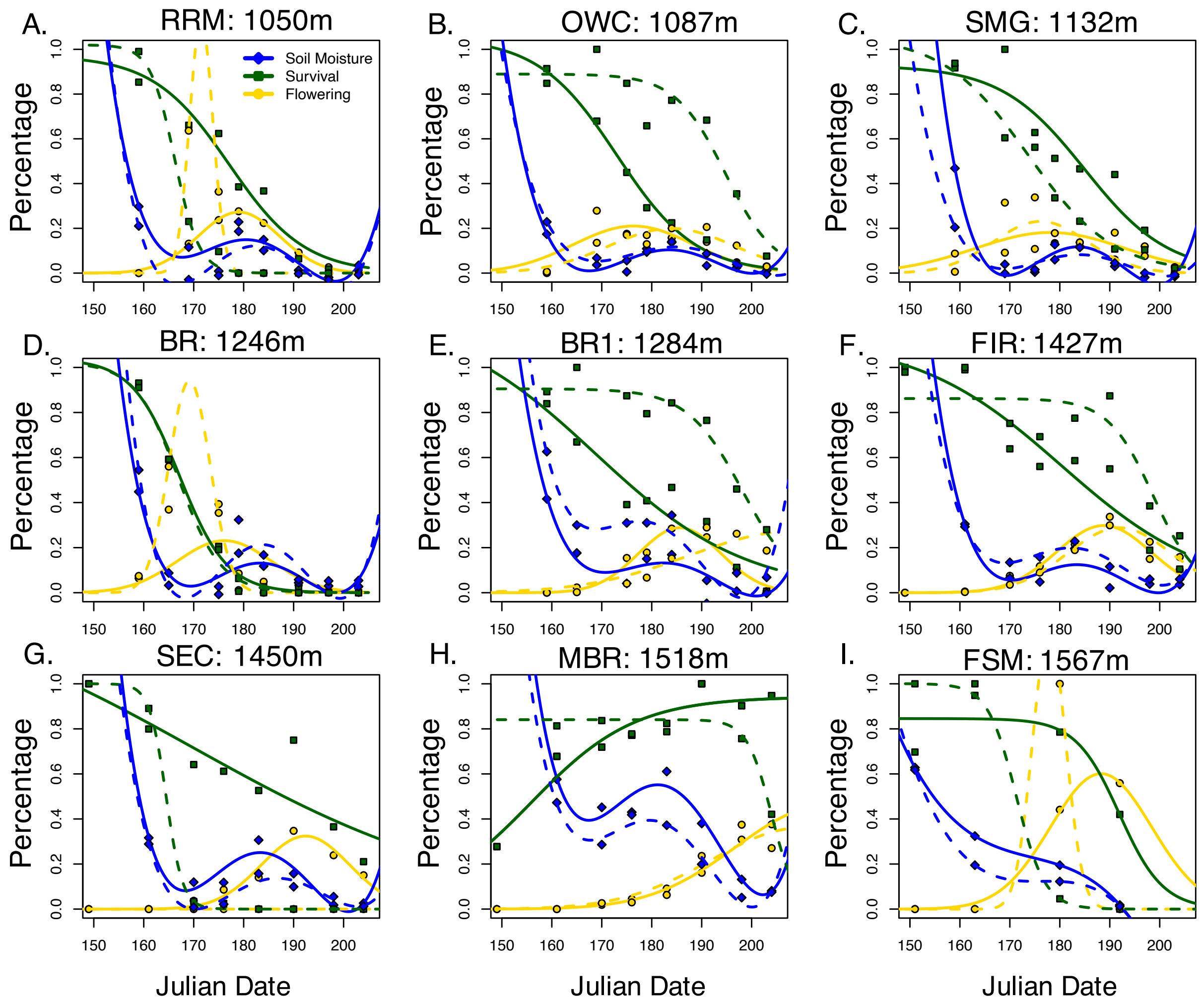


**Fig. S4** Phenology during the 2019 growing season for Browder Ridge monkeyflower populations. Each scatterplot (A-I) depicts volumetric water content (blue), proportion of the population surviving (green), and proportion of the population flowering (yellow) during the growing season for a single population. Three letter abbreviations and elevations are given above each graph. Lines represent inferred models for each of the two grids within a population and different grids are represented by dashed vs solid lines.

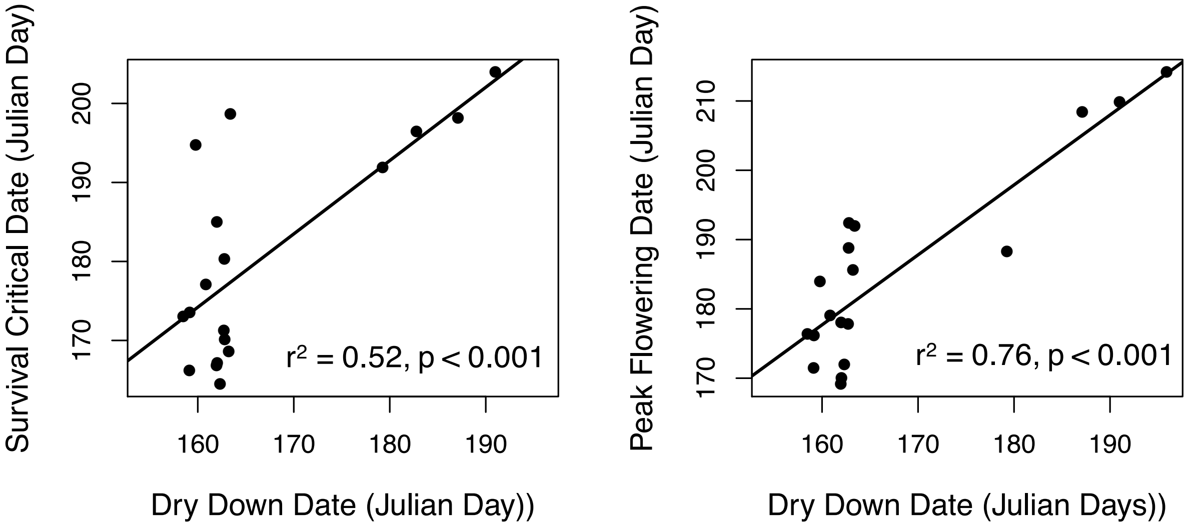


**Fig. S5** Relationships between dry down dates and population phenology for Browder Ridge monkeyflower populations. Scatterplots depict associations between dry down date and either critical survival date (A) or flowering peak date (B). Dry down date was defined as the predicted day where volumetric water content drops below 0.2. Critical survival date was defined as the inflection point on a survival time series model. Peak flowering date was defined as the predicted day of peak flowering from fitted models.
