## Supplemental Methods for "Demographic consequences of an extreme heatwave are mitigated by spatial heterogeneity in an annual monkeyflower"

These methods describe how we modeled mortality, phenology and soil moisture through growing season and calculating summary statistics for each grid within each population. We assessed the proportion of plants surviving at a given time point by dividing the number of plants alive by the maximum census within the sampling grid. At some high elevation sites, we started censusing right after snowmelt when there were no germinates at first sampling, thus the maximum census size occurred at the second or third sampling point. To examine how survival curves differed within and among populations we used non-linear least square regression to fit logistic curves implemented with the *SSlogis()* function from the *stats* v3.6.2 package. To compare between grids and populations we used this model to estimate when 50% survival occurred during the growing season (termed critical survival date below). We also estimated the end of the growing season by calculating when survivorship was 5%. To assess flowering phenology, we divided the number of plants flowering in a grid at a given date by the total number of plants that had flowered throughout the experiment within that sampling grid. This measure is flawed as the same plants may be recounted as flowering multiple times, however the relative phenology comparison between plots is still informative for our purposes. To compare within and between populations, we fit gaussian curves to phenology data and optimized the fit via the *optim()* function to minimize the sum of squared residuals. We used this model to estimate when flowering began in each grid (i.e., when 5% of plants had flowered) and when flowering peaked in each plot. To model how soil moisture changed throughout the growing season we fit fourth-order polynomial models using the *lm()* function. The fourth order model was selected because it best accounted for late season increases in soil moisture that improved fitness in some populations. To compare patterns of dry down between populations, we calculated the Julian day when volumetric water content (VWC) first reached 0.2 using fitted polynomial models. This value was selected based on previous dry-down experiments with *M. guttatus* in controlled conditions (N. Kooyers pers. obs).
