## Supplemental Tables for "Demographic consequences of an extreme heatwave are mitigated by spatial heterogeneity in an annual monkeyflower"

**Table S1.** Summary of ANOVA results examining effects of population on survival within the common garden.

| Response Variable | Model: Family (Link) | Fixed Factor | $X^2$ | df | p |
|---|---|---|---|---|---|
| Survival to Flowering | GLMM: Binomial (Logit) | Population | 22.9 | 10 | 0.011 |
| Inclusive Number of Flowers | LMM | Population | 47.0 | 10 | < 0.001 |
| Inclusive Number of Seeds | LMM | Population | 40.1 | 10 | < 0.001 |
| Inclusive Number of Flowers | GLMM:Poisson (Logit) | Population | 41.4 | 10 | < 0.001 |
| Inclusive Number of Seeds | GLMM:Poisson (Logit) | Population | 20.5 | 10 | 0.025 |

Inclusive number of flowers of flowers or seeds refers to number of flowers or seeds with anyone individual producing no flowers or seeds having a value of zero. Inclusive number of seeds was log+1 transformed for all LMM. LMM = Linear Mixed Model; GLMM = Generalized Linear Mixed Model

**Table S2.** Fitness summary statistics for populations within the common garden experiment

| Pop. | N | Lat. | Elev. (m) | Dist. (km) | AHM | Survival to Flower | Flowers Ave. (SD) | Seeds Ave. (SD) | Inc. Number of Flowers Ave. (SD) | Inc. Number of Seeds Ave. (SD) |
|---|---|---|---|---|---|---|---|---|---|---|
| BR1 | 8 | 44.37 | 1269 | 0.4 | 6.6 | 0 (0) | NA | NA | 0 (0) | 0 (0) |
| MTC | 5 | 44.23 | 1400 | 15.6 | 7.7 | 0.03 (0.07) | 1.5 (NA) | 3 (NA) | 0.05 (0.11) | 0.1 (0.22) |
| LPD | 8 | 43.92 | 277 | 71.2 | 17.6 | 0.12 (0.1) | 1.67 (0.51) | 42 (39) | 0.2 (0.2) | 4.94 (6.94) |
| SWC | 8 | 44.01 | 204 | 143.6 | 9.9 | 0.02 (0.04) | NA | NA | 0.02 (0.04) | 0 (0) |
| GBS | 6 | 42.42 | 93 | 284.3 | 10.3 | 0 (0) | NA | NA | 0 (0) | 0 (0) |
| TAR | 6 | 40.85 | 778 | 396.1 | 22.8 | 0.18 (0.14) | 1.06 (0.83) | 25 (12) | 0.22 (0.18) | 2.51 (2.87) |
| BLD | 5 | 38.14 | 1733 | 714.1 | 18.7 | 0.05 (0.11) | 2 (NA) | 133 (NA) | 0.08 (0.18) | 3.99 (8.92) |
| YVO | 8 | 37.72 | 1495 | 765.9 | 22.1 | 0.04 (0.07) | 1.5(NA) | 64 (NA) | 0.05 (0.09) | 0.67 (1.89) |
| BEL | 8 | 37.04 | 196 | 839.2 | 75.9 | 0.19 (0.16) | 2.08 (0.74) | 95 (75) | 0.39 (0.4) | 17.55 (23.9) |
| WPA | 8 | 35.43 | 377 | 1003.8 | 41.7 | 0.06 (0.06) | 1 (0) | 13 (13) | 0.06 (0.06) | 0.31 (0.77) |
| MUG | 8 | 32.96 | 331 | 1346.9 | 71.3 | 0.02 (0.05) | NA | NA | 0.02 (0.05) | 0 (0) |

Fitness measures are all given in the format: mean (standard deviation). Inc. number of flowers or seeds refer to the number of flowers or seeds respectively and include all plants that did not survive to flowering. Abbreviations: Pop. = Population, N = Number of maternal lines per population, Elev. = elevation, Dist. = Distance from the common garden site, AHM = Annual Heat Moisture Index

**Table S3.** Model results examining environmental associations with fitness metrics for populations within the common garden.

| Response Variable | Model | Independent Variable | $X^2$ | df | P-Values |
|---|---|---|---|---|---|
| Distance to Common Garden | GLMM: Binomial (Logit) | Survival to Flowering | 0.0135 | 1 | 0.9075 |
| Annual Heat Moisture Index | GLMM: Binomial (Logit) | Survival to Flowering | 0.4303 | 1 | 0.5119 |
| Mean Annual Temperature | GLMM: Binomial (Logit) | Survival to Flowering | 0.202 | 1 | 0.6531 |
| Beginning of the Frost Free Period | GLMM: Binomial (Logit) | Survival to Flowering | 0.0621 | 1 | 0.8032 |
| Climate Moisture Deficit | GLMM: Binomial (Logit) | Survival to Flowering | 0.9858 | 1 | 0.3208 |
| Variance in Spring Max Temperature | GLMM: Binomial (Logit) | Survival to Flowering | 3.5907 | 1 | 0.0581 |
| Variance in Summer Max Temperature | GLMM: Binomial (Logit) | Survival to Flowering | 0.8803 | 1 | 0.3481 |
| Distance to Common Garden | LMM | Inclusive Number of Flowers | 0.0022 | 1 | 0.9629 |
| Annual Heat Moisture Index | LMM | Inclusive Number of Flowers | 1.1446 | 1 | 0.2847 |
| Mean Annual Temperature | LMM | Inclusive Number of Flowers | 0.5169 | 1 | 0.4722 |
| Beginning of the Frost Free Period | LMM | Inclusive Number of Flowers | 0.1221 | 1 | 0.7268 |
| Climate Moisture Deficit | LMM | Inclusive Number of Flowers | 0.7655 | 1 | 0.3816 |
| Variance in Spring Max Temperature | LMM | Inclusive Number of Flowers | 1.2113 | 1 | 0.2711 |
| Variance in Summer Max Temperature | LMM | Inclusive Number of Flowers | 0.205 | 1 | 0.6508 |
| Distance to Common Garden | LMM | Inclusive Number of Seeds | 0.0403 | 1 | 0.841 |
| Annual Heat Moisture Index | LMM | Inclusive Number of Seeds | 1.7181 | 1 | 0.1899 |
| Mean Annual Temperature | LMM | Inclusive Number of Seeds | 0.8435 | 1 | 0.3584 |
| Beginning of the Frost Free Period | LMM | Inclusive Number of Seeds | 0.2657 | 1 | 0.6062 |
| Climate Moisture Deficit | LMM | Inclusive Number of Seeds | 1.1512 | 1 | 0.2833 |
| Variance in Spring Max Temperature | LMM | Inclusive Number of Seeds | 1.3747 | 1 | 0.241 |
| Variance in Summer Max Temperature | LMM | Inclusive Number of Seeds | 0.1126 | 1 | 0.7372 |

Inclusive number of flowers or seeds refer to the number of flowers or seeds respectively and include all plants that did not survive to flowering. Female fitness was log+1 transformed for all models. LMM = Linear Mixed Model; GLMM = Generalized Linear Mixed Model

**Table S4.** Mortality and phenology of natural populations near the common garden site.

| Site | Elev. (m) | Seeds 2018 Ave.(SD) | Seeds 2019 Ave.(SD) | Flowers 2018 Ave.(SD) | Flowers 2019 Ave.(SD) | Grid | Alive (initial) | Alive (before) | Alive (after) | Mortality Rate | VWC Date (<0.2) | Surv. Date (50% Surv.) | Flowering Peak Date |
|---|---|---|---|---|---|---|---|---|---|---|---|---|---|
| RRM | 1050 | 124.3 (176.9) | 41.8 (64.9) | 5.2 (4.3) | 2.4 (1.7) | A | 109 | 93 | 68 | 0.27 | 160.8 | 177.1 | 179 |
| | | | | | | B | 104 | 103 | 10 | 0.9 | 159.1 | 166.2 | 171.5 |
| OWC | 1087 | 55.4 (88.6) | 68.4 (104.5) | 3 (2.1) | 2.4 (1.7) | A | 209 | 191 | 94 | 0.51 | 158.5 | 173.1 | 176.4 |
| | | | | | | B | 79 | 67 | 67 | 0 | 159.8 | 194.8 | 184 |
| SMG | 1132 | 62.5 (73.1) | 74.4 (130.3) | 4.6 (3.3) | 4.1 (4.2) | A | 231 | 254 | 174 | 0.32 | 162 | 185 | 178.1 |
| | | | | | | B | 354 | 332 | 199 | 0.4 | 159.2 | 173.6 | 176.1 |
| BR | 1246 | - | - | - | - | A | 794 | 739 | 163 | 0.78 | 162 | 167.2 | 170.1 |
| | | | | | | B | 324 | 295 | 62 | 0.79 | 162 | 166.9 | 169.2 |
| BR1 | 1284 | - | - | - | - | A | 1051 | 882 | 411 | 0.53 | 163.2 | 168.6 | 185.6 |
| | | | | | | B | 537 | 539 | 528 | 0.02 | 187.1 | 198.2 | 208.4 |
| FIR | 1427 | 13.7 (19.7) | 31.9 (36) | 1.6 (1.1) | 1.8 (0.9) | A | 191 | 189 | 122 | 0.35 | 162.8 | 180.3 | 188.8 |
| | | | | | | B | 465 | 475 | 357 | 0.25 | 163.4 | 198.7 | 191.9 |
| SEC | 1450 | 23.1 (28.3) | 64.3 (106.7) | 1.8 (1.2) | 2.5 (2.1) | A | 231 | 254 | 174 | 0.32 | 162.8 | 170.1 | 192.4 |
| | | | | | | B | 119 | 106 | 3 | 0.97 | 162.3 | 164.5 | 172 |
| HDM | 1465 | 24.6 (31.1) | - | 2.5 (1.7) | - | A | 124 | 50 | 0 | 1 | 156.3 | - | - |
| | | | | | | B | 12 | 2 | 0 | 1 | 154.4 | - | - |
| FSM | 1535 | 51.2 (59.4) | 10.5 (20.7) | 2.4 (1.8) | 1.6 (0.8) | A | 244 | 350 | 275 | 0.21 | 179.3 | 191.9 | 188.3 |
| | | | | | | B | 175 | 166 | 8 | 0.95 | 162.7 | 171.3 | 177.9 |
| MBR | 1518 | 41.2 (53.7) | 13.1 (19.8) | 2.8 (2.7) | 1.3 (0.7) | A | 209 | 191 | 94 | 0.51 | 195.9 | - | 214.2 |
| | | | | | | B | 66 | 344 | 329 | -0.04 | 191 | 204 | 209.9 |
| HOV | 1567 | 64.7 (67.3) | 42.9 (55) | 3.9 (2.8) | 2.2 (1.2) | A | 183 | 187 | 178 | 0.05 | 182.8 | - | - |
| | | | | | | B | 81 | 137 | 1 | 0.99 | 165.2 | - | - |

Dashes indicate that models were not run because either there was no survival following the heatwave or because phenology surveys ended before substantial decline of plants and flowering within a grid. All dates are Julian dates.

**Table S5.** Associations between phenology and environment variables within natural populations.

| Response Variable | Independent Variable | Sum of Squares | Mean Square | Numerator DF | Denominator DF | F-value | P-value |
|---|---|---|---|---|---|---|---|
| Dry Down Date | Elevation | 154.9 | 154.9 | 1 | 9 | 2.85 | 0.126 |
| Dry Down Date | MAT | 251.3 | 251.3 | 1 | 8.9966 | 4.63 | 0.06 |
| Dry Down Date | MCMT | 330.4 | 330.4 | 1 | 9 | 6.08 | 0.036 |
| Dry Down Date | bFFP | 45.1 | 45.1 | 1 | 9 | 0.83 | 0.386 |
| Dry Down Date | PAS | 267 | 267 | 1 | 9 | 4.92 | 0.054 |
| Critical Survival Date | Elevation | 213.4 | 213.4 | 1 | 7.4632 | 1.36 | 0.279 |
| Critical Survival Date | MAT | 118.8 | 118.8 | 1 | 7.4655 | 0.75 | 0.412 |
| Critical Survival Date | MCMT | 139.6 | 139.6 | 1 | 7.871 | 0.89 | 0.372 |
| Critical Survival Date | bFFP | 22.1 | 22.1 | 1 | 5.6088 | 0.13 | 0.729 |
| Critical Survival Date | PAS | 173.5 | 173.5 | 1 | 8.2698 | 1.12 | 0.32 |
| Flowering Peak | Elevation | 219.9 | 219.9 | 1 | 7 | 3.32 | 0.111 |
| Flowering Peak | MAT | 206.6 | 206.6 | 1 | 7 | 3.12 | 0.121 |
| Flowering Peak | MCMT | 192.6 | 192.6 | 1 | 7 | 2.91 | 0.132 |
| Flowering Peak | bFFP | 10.6 | 10.6 | 1 | 7 | 0.16 | 0.701 |
| Flowering Peak | PAS | 212.3 | 212.3 | 1 | 7 | 3.21 | 0.117 |

Dry down date refers to the date that a grid registered a VWC of < 0.2. Critical Survival Date is the Julian day of the year with 50% of plants remaining. Flowering peak is the model estimate of Julian date with the highest frequency of plants flowering. Each response variable was derived from the corresponding models rather than actual observation dates. Associations between response variables and environmental predictor variables were determined with linear mixed models as described in the text. Abbreviations: MAT= Mean Annual Temperature, MCMT = Mean Coldest Month Temperature, bFFP = Beginning of the Frost Free Period, PAS = Precipitation as Snow.

**Table S6.** Associations between natural population fecundity in 2018 and 2019 with environmental characteristics

| Response Variable | Environmental Predictor | r2 | F-Value | Degrees of Freedom | P-Value |
|---|---|---|---|---|---|
| Difference in Average Seed Production | Elevation | 0.04 | 0.38 | 1,10 | 0.55 |
| Difference in Average Seed Production | MAT | 0.03 | 0.35 | 1,10 | 0.57 |
| Difference in Average Seed Production | MCMT | 0.03 | 0.35 | 1,10 | 0.57 |
| Difference in Average Seed Production | bFFP | 0.03 | 0.26 | 1,10 | 0.62 |
| Difference in Average Seed Production | PAS | 0.02 | 0.24 | 1,10 | 0.64 |
| Difference in Average Seed Production | CMD | 0.11 | 1.28 | 1,10 | 0.28 |
| Difference in Average Flower Production | Elevation | 0.09 | 1.03 | 1,10 | 0.34 |
| Difference in Average Flower Production | MAT | 0.06 | 0.64 | 1,10 | 0.44 |
| Difference in Average Flower Production | MCMT | 0.03 | 0.34 | 1,10 | 0.57 |
| Difference in Average Flower Production | bFFP | 0.05 | 0.49 | 1,10 | 0.5 |
| Difference in Average Flower Production | PAS | 0.01 | 0.12 | 1,10 | 0.74 |
| Difference in Average Flower Production | CMD | 0.16 | 1.85 | 1,10 | 0.2 |

Differences in seed and flower production between 2018 and 2019 were calculated from population averages. Abbreviations: MAT= Mean Annual Temperature, MCMT = Mean Coldest Month Temperature, bFFP = Beginning of the Frost Free Period, PAS = Precipitation as Snow, CMD = Climate Moisture Deficit
